# Case study: Paralog diverged features may help reduce off-target effects of drugs

**DOI:** 10.1101/078063

**Authors:** Zhining Sa, Jingqi Zhou, Yangyun Zou, Xun Gu

**Author notes:** Corresponding author: Xun Gu,; Yangyun Zou. These authors contributed equally to this work and should be considered co-first authors (Zhining Sa, Jingqi Zhou).

## Abstract

Side effects from targeted drugs is a serious concern. One reason is the nonselective binding of a drug to unintended proteins such as its paralogs, which are highly homologous in sequences and exhibit similar structures and drug-binding pockets. In this study, we analyzed amino acid residues with type-II functional divergence, i.e., sites that are conserved in sequence constraints but differ in physicochemical properties between paralogs, to identify targetable differences between two paralogs. We analyzed paralogous protein receptors in the glucagon-like subfamily, glucagon receptor (GCGR) and glucagon-like peptide-1 receptor (GLP-1R), which are clinically validated drug targets in patients with type 2 diabetes and exhibit divergence in ligands, showing opposing roles in regulating glucose homeostasis. We identified 8 residues related to type-II functional divergence, which are conserved in functional constraints but differ in physicochemical properties between GCGR and GLP-1R. We detected significant enrichment of predicted residues in binding sites of the antagonist MK-0893 to GCGR. We also identified a type-II functional divergence-related residue involved in ligand-specific effects that was critical for agonist-mediated activation of GLP-1R. We describe the important role of type-II functional divergence-related sites in paralog discrimination, enabling the identification of binding sites to reduce undesirable side effects and increase the target specificity of drugs.

## Introduction

Precision medicine, as an emerging area and therapeutic strategy [1], enables the development of targeted drugs and improves the efficacy of therapy. However, some targeted drugs are promiscuous, showing a high risk of severe side effects because they have unexpected targets and exhibit low specificity [2]. Cross-reactivity on protein paralogs may cause undesirable side effects of drugs [3]. Paralogs are evolutionarily homologous and are generated from duplications [4]. They share similar protein sequences or structural features, thus comprising similar binding pockets with drugs. A drug that binds to one gene target may also bind to its paralog, often resulting in unexpected cross-reactivity and leading to undesired side effects. Therefore, rationally controlling specificity to limit side effects is required to create novel and safer drugs. This control may be achieved by drug design guided by paralog-discriminating features, known as “selectivity filters” [3]. One strategy for achieving specificity is identifying evolutionarily divergent features that enable paralog discrimination. This method is based on the association between the change in the evolutionary rate and functional divergence after gene duplication by applying the underlying fundamental rule that amino acids are evolutionarily conserved if they are functionally important [5]. A shift in key physicochemical properties relevant to ligand binding interactions may result in changes in binding features or considerably affect the druggability of protein targets [6]. Type-II functional divergence-related sites refer to residues that are evolutionarily conserved but differ in physicochemical properties, e.g., positive versus negative charge differences between paralogous sites, which are typically known as “constant but different” [7,8]. Therefore, these divergent features in physicochemical properties between paralogs can be exploited as selectivity filters to function as targetable differences [9].

In this study, we investigated the known target protein receptor family G-protein coupled receptors (GPCRs), which highly contribute to side effects [10]. We used glucagon-like subfamily of secretin-type GPCRs as an example to illustrate our analytical pipeline for detecting paralog diverged features, e.g. type-II functional divergence sites between duplicate clusters. These features can be considered in the drug design of known drugs such as the GCGR antagonist MK-0893. We also describe the important role of type-II functional divergence between GCGR and GLP-1R in paralog discrimination, which may be useful for identifying binding sites to achieve target specificity and develop safer and more selective drugs.

## Materials and Methods

### Data sets

We retrieved 319 unique functional nonolfactory human GPCRs from the *GRAFS* classification system previously proposed by Fredriksson [11]. Druggable genes belonging to an orthologous quartet (derived from the human, macaque, mouse, and rat genomes) were obtained from a previous study [12]. We identified druggable GPCRs from these 1,362 genes with additional published [13,14] data. Finally, we identified 82 G-protein coupled receptors as drug targets.

### Multiple alignment and phylogenetic analysis

We downloaded 41 amino acid sequences of the glucagon-like subfamily in human GPCRs as well as their vertebrate and invertebrate orthologs from the ENSEMBL database. To maintain uniqueness, partial and redundant sequences were removed, and only those genes with the longest proteins sequences were retained for further analysis. The multiple alignment of amino acid sequences was conducted using MEGA 7.0 software [15]. Gaps were removed, and a phylogenetic tree of glucagon-like subfamily was inferred by the neighbor-joining method with Poisson distance. Similar results were obtained using other methods (i.e. parsimony, maximum likelihood, and Bayesian methods; results not shown). The concordance of the results from different phylogenetic methods increased the confidence in the relationships inferred from the presented tree. A phylogenetic tree of GCGR and GLP-1R was constructed in the same manner.

### Analytical pipeline for type-II functional divergence analysis

We used DIVERGE3.0 [16] to explore the functional evolution of glucagon-like subfamily sequences. The site-specific profiles of two duplicate genes clusters were determined to detect amino acid residues that are crucial for type-II functional divergence. A typical case is that site-specific property shifts between duplicate genes, e.g., positively vs. negatively charged, but is highly conserved within the cluster. Amino acids are classified into four groups [17]: charge positive (K, R, H), charge negative (D, E), hydrophilic (S, T, N, Q, C, G, P), and hydrophobic (A, I, L, M, F, W, V, Y). When an amino acid changes from one group to another, it is referred to as radical; otherwise, it is conserved. We used the coefficient θ_II_ to measure the level of type-II functional divergence between two clusters. A larger θ_II_ implies a stronger type-II functional divergence. Thus, we first tested whether θ_II_ > 0. Next, we determined the posterior ratio R_II_ (k) = Q_II_ (k)/ [1- Q_II_ (k)], where Q_II_ (k) is a site (k)-specific score. Amino acid residues with radical changes between duplicate clusters received higher scores than those with conserved changes in physicochemical properties. Under a given cut-off value, we screened important residues related to type-II functional divergence between duplicated genes.

### Schematic topological representation

We used snake-plot diagrams produced using web tools in the GPCRdb database [18] to illustrate receptor residues of interest.

### PDB structure

We downloaded the crystal structure of human glucagon receptor (GCGR) in complex with the antagonist MK-0893, the chain A of the PDB ID 5EE7 from RCSB Protein database [19]. Next, we utilized PyMOL software [20] to illustrate the mechanism of target binding and clarify the relationship of type-II residues with antagonist binding sites.

## Results

### Case study: glucagon-like subfamily

GPCRs constitute one of the largest families of membrane proteins with approximately 800 members in the human genome [21]. It is estimated that 30–40% of all drugs currently on the market target GPCRs [22] (Figure 1a). The glucagon-like subgroup is one of the subfamilies in secretin-type GPCRs, which is rich in clinically validated targets [23]. This family constitutes 4 hormone receptors duplicated from the early stage of vertebrates [24] (Figure 1b). These receptors play a crucial role in hormonal homeostasis in humans and other animals and serve as important drug targets for several endocrine disorders [25]. Among them, the glucagon receptor (GCGR) and glucagon-like peptide-1 receptor (GLP-1R) appear to have greater therapeutic potential in diabetes than the other members [24,26,27]. Thus, we focused on GCGR and GLP-1R for further investigation.

**Figure 1.**
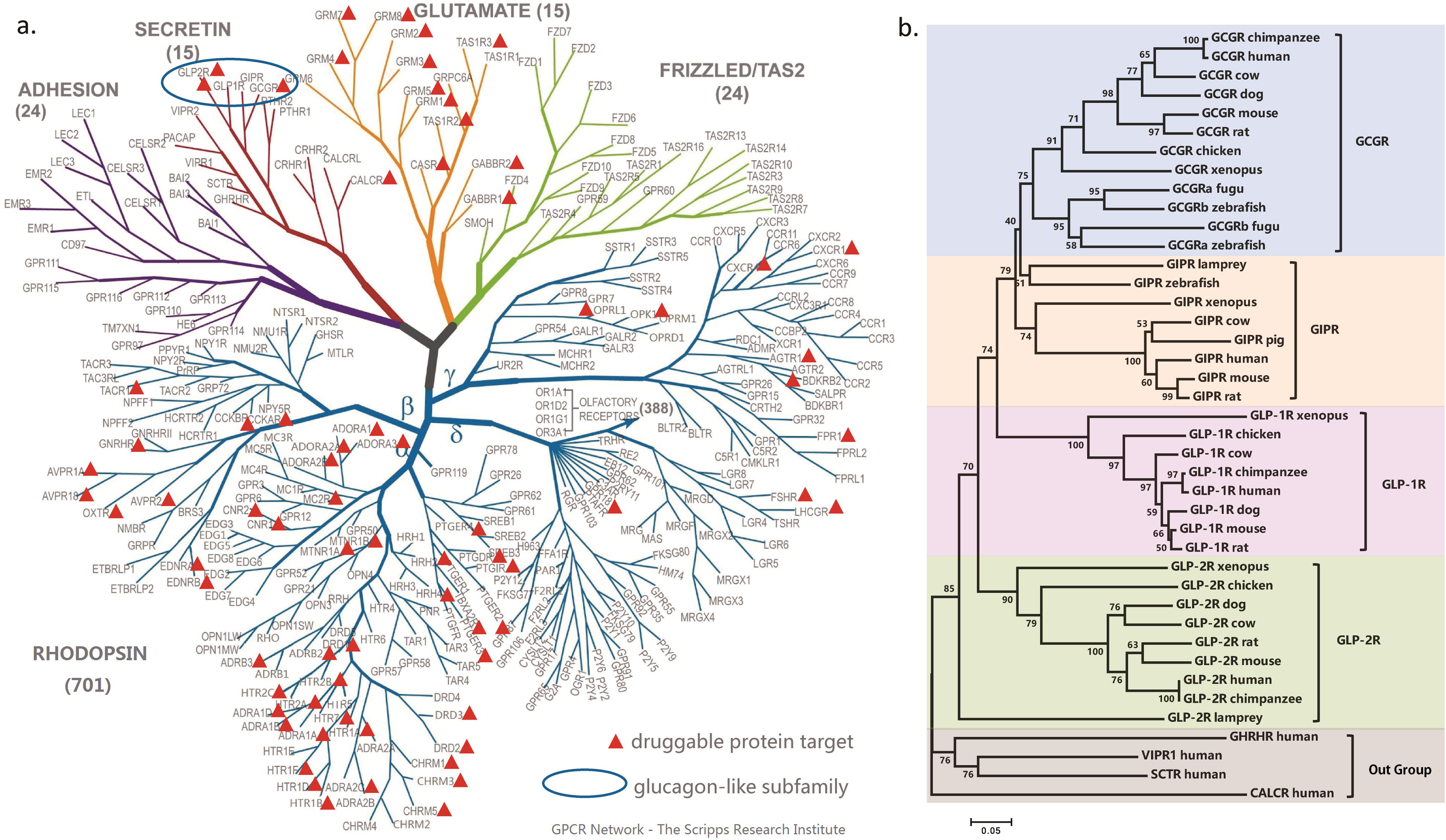
GPCRs are likely to be drug targets. a) 82 targetable receptors are plotted on the GPCRs tree (courtesy of V. Katritchb and R. C. Stevens - Scripps/USC). b) Lineage divergence of drug targets in glucagon-like subfamily.

GCGR shares high homology with GLP-1R, where the sequence identities in the transmembrane and extracellular domains are, respectively, 54.0% and 46% [28,29]. In addition, the corresponding ligands for GCGR and GLP-1R, glucagon and GLP-1, respectively, are also highly conserved in sequence [30]. It has been hypothesized that GLP-1R exhibits glucagon-like action in fish in the early stage, but later acquires unique incretin functions [31]. In mammals, these two hormones have significant but opposing roles in regulating glucose homeostasis and are clinically important in the management of diabetes [32]. Glucagon acts primarily on hepatic GCGR to increase plasma glucose, while GLP-1 functions during nutrient ingestion at pancreatic β-cell GLP-1R to enhance insulin synthesis and secretion [29]. GLP-1 affects blood glucose, β-cell protection, appetite, and body weight, which has led to the use of multiple GLP-1R agonists for the treatment of type 2 diabetes [33]. In contrast, glucagon is used to treat severe hypoglycemia [34], while antagonists have been developed to treat type 2 diabetes. Thus, GCGR and GLP-1R show divergent ligand binding profiles and are selective in hormone action, although they are highly homologous and show conserved structures and sequences. Therefore, when GCGR antagonists wrongly target highly homologous GLP-1R in patients with type 2 diabetes, these drugs may lose their efficacy by not controlling the release of glucose by GCGR and influence the function of GLP-1R by decreasing the augmentation of insulin secretion. As a result, anti-diabetes drugs targeting one of these two paralogous receptors at conserved sites may also target the other one by mistake, resulting in cross-reactivity and generating unexpected side effects.

### Identification of paralog diverged features among glucagon-like subfamily

To avoid undesirable side effects driven by drug interactions with conserved residues of paralogs, we analyzed type-II functional divergence between GCGR and GLP-1R to identify residues conserved in functional constraints but differing in physicochemical properties. Based on the phylogenetic tree and sequence configuration (Figure 2a), we estimated the coefficient of type-II functional divergence (denoted by θ_II_) between GCGR and GLP-1R, θ_II_ = 0.236 ± 0.052, which showed a value significantly larger than 0 (p-value < 0.001). This suggests that after gene duplication, some amino acid residues that were evolutionarily conserved in both GCGR and GLP-1R may have radically changed their amino acid properties. Further, we used the posterior ratio RII (k) to identify amino acid residues critical in type-II functional divergence between these two paralogous genes (Figure 2b). We used an empirical cutoff of R_II_ (k) > 2 (posterior probability Q_II_ (k) > 0.67) to identify 8 type-II functional divergence-related residues (Glu34, Ser150, Asn291, Gln337, Phe345, Phe387, Lys405, and Glu427 in GCGR) between paralogous GCGR and GLP-1R. The site-specific ratio profile indicated that most residues had low posterior ratios and only a small portion of amino acid residues were involved in this type of functional divergence. Moreover, these 8 amino acid residues showed a typical pattern of type-II functional divergence (Figure 2c). They showed sequence conservation and functional constraints at paralogous sites (Figure 2d), while the lower cut-off value led to variable amino acid residues in both paralogs. Thus, we used these 8 type-II functional-specific sites for further analysis to gain insights into their roles in paralog discrimination.

**Figure 2.**
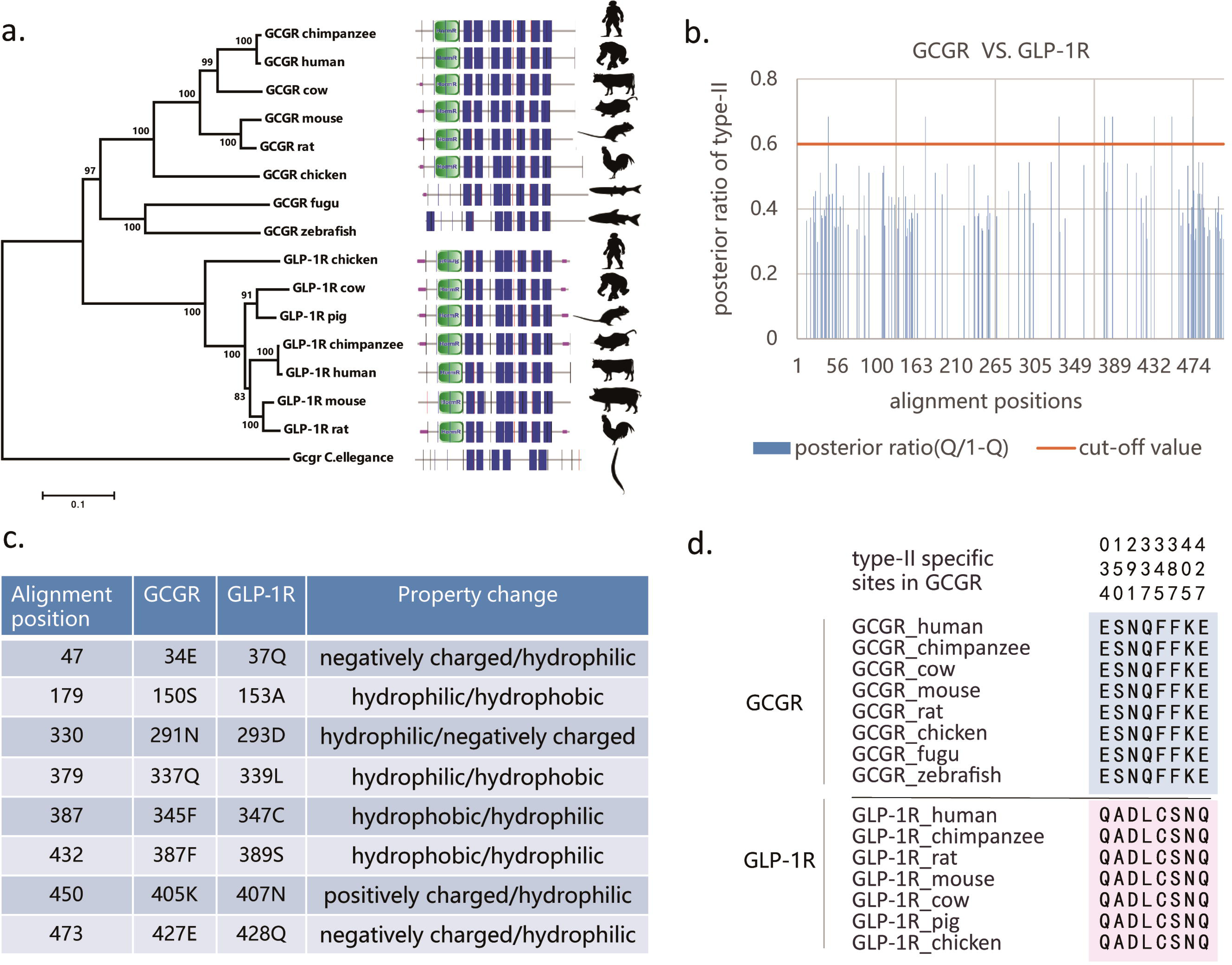
Analytical pipeline for type-II functional divergence between GCGR and GLP-1R. a) Phylogenetic tree of GCGR and GLP-1R with domain information. b) Site-specific profile for predicting critical amino acid residues responsible for type-II functional divergence between GCGR and GLP-1R measured by posterior ratio R_II_ (k). c) Overview of amino acid changes in the 8 predicted sites in type-II functional divergence. d) Sequence conservation analysis of two clusters.

### Usage of paralog diverged features as targetable difference of drugs

The cross-reactivity arising from paralogs is considered to be one cause of the side effects of drugs. Because most drug targets are paralogs, a method for identifying targetable differences is necessary for the design of therapeutic drugs. We hypothesized that paralog diverged features such as type-II functional divergence between two duplicated clusters may be a possible solution. The GCGR antagonist MK-0893 is used to treat patients with type 2 diabetes to substantially reduce fasting and postprandial glucose concentrations [35]. MK-0893 acts at allosteric binding sites of the seven transmembrane helical domain (7TM) in positions among TM5, TM6, and TM7 in GCGR (Figure 3a). TM6 plays a role in dividing the binding sites into two different interaction regions. The TM5-TM6 cleft includes Leu329, Phe345, Leu352, Thr353, and the alkyl chain of Lys349, which makes hydrophobic contacts with one part of MK-0893. The TM6-TM7 section forms polar interactions with another part of MK-0893 by hydrogen bonding with Lys349, Ser350, Leu399, Asn404, and the backbone of Lys405, and additional salt bridge with Arg346. Thus, the different physicochemical properties function in the binding activity of the dual-nature antagonist MK-0893 to GCGR (Figure 3b). We found that our predicted sites of type-II functional divergence between GCGR and GLP-1R. Phe345 and Lys405 were significantly enriched in the binding sites of MK-0893 to GCGR (chi-square test is statistically significant with p-value < 0.05). Further, we compared these allosteric sites with equivalent sites in GLP-1R. The results showed that these binding sites were highly conserved either in functional constraints or physicochemical properties between two paralogs except for the type-II-specific sites Phe345 and Lys405 (Figure 3c). Phe345, showing a typical pattern of type-II functional divergence, was hydrophobic in GCGR and hydrophilic in GLP-1R. Another type-II site Lys405 was positively charged in GCGR and was hydrophilic in GLP-1R. Because the physiochemical properties of amino acids play an important role in the interaction of protein receptors with their ligands (small molecules, peptides, agonists, and antagonists), changes in their physicochemical nature and conformation may reduce cross-reactivity resulting from antagonist pockets binding to unexpected paralogs. Therefore, determining type-II functional divergence-related sites between two paralogs is effective for identifying targetable differences in therapeutic drug design.

**Figure 3.**
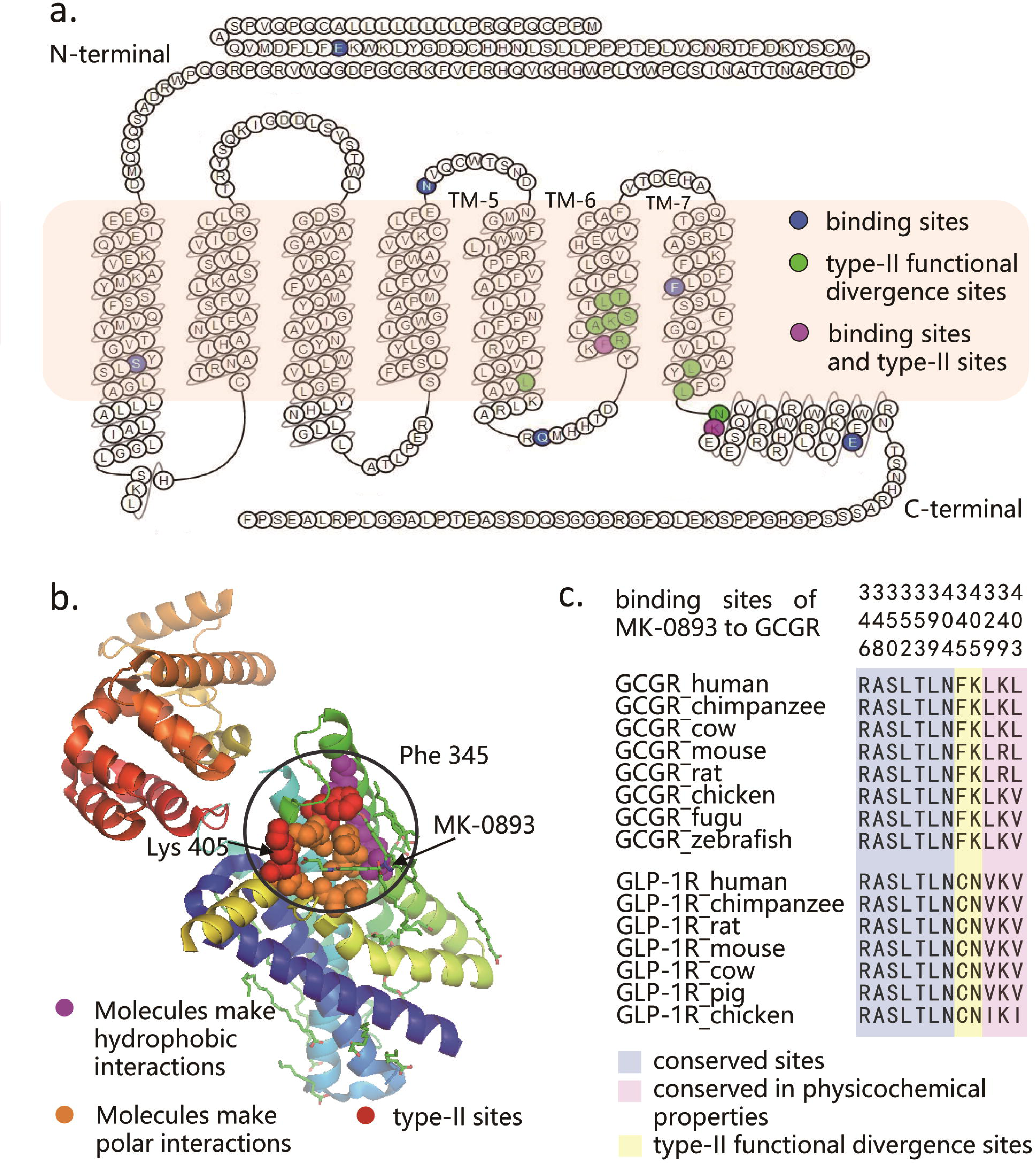
Paralog diverged features are considered targetable differences of drugs. a) Snake-plot diagram of GCGR with annotation of important residues. b) Different physicochemical properties of bipartite antagonist pocket corresponding to the dual polar/hydrophobic nature of binding cleft in GCGR. c) Sequence conservation analysis of 12 binding sites of MK-0893 to GCGR

Moreover, we investigated the binding of ligand and agonists GLP-1R and evaluated the role of type-II functional divergence sites between GCGR and GLP-1R in this study. We identified a type-II functional divergence-related residue Asp293 within human GLP-1R in the second extracellular loop (EC2), which had ligand-specific effects on GLP-1 peptide-mediated selective signaling and was critical for agonist-mediated receptor activation [36]. Residue Asp293 of EC2 directly interacted with key residues in the ligand through hydrogen-bonding interactions. A previous study [37] demonstrated that a mutation in this residue to alanine reduced GLP-1 affinity and altered the binding and efficacy of agonists such as oxyntomodulin and exendin-4 [38]. A functionally important site such as Asp293 showed sequence conservation but different physicochemical properties of the amino acid between paralogous GLP-1R and GCGR. Thus, the application of divergent features of type-II functional divergence between these two paralogs is advantageous in this respect. The amino acid property changes from negatively charged in GLP-1R to hydrophobic in GCGR can serve as a selective filter for differentiating GLP-1R and GCGR.

### Discussion

The side effects of drugs arise from off-target effects (nonselective binding to other proteins besides the intended targets) [39]. Because paralogous proteins share similar structures and sequences, a drug that targets one paralog is likely to bind to other paralogs as well [40]. Although affinity toward these paralogs can be lower than to the intended protein targets, the number of off-target paralogs can be sufficiently high to mediate the side effects [41]. Therefore, side effects due to paralogous binding must be controlled and target selectivity improved for rational drug design. Here, we used an analytical pipeline to determine the type-II functional divergence between GCGR and GLP-1R to identify residues that can be regarded as targetable differences. We used the antagonist MK0893 to target GCGR and found that our predicted type-II functional divergence-related residues were significantly enriched in the binding sites of GCGR. The type-II residues Phe345 and Lys405 showed a radical shift in physicochemical properties between GCGR and GLP-1R, while other binding sites were highly conserved between the two paralogs. Thus, type-II functional divergence-related sites may be critical in paralog discrimination. Undesirable side effects can occur because GCGR and GLP-1R have diverged in ligands and exhibit very different roles in regulating glucose homeostasis. Further, we observed another type-II residue, Asp293, in the binding sites of GLP-1R interacting with residues of its ligand and agonists. Asp293 is a functionally important site and its variation in physicochemical properties can differentiate paralogous GCGR and GLP-1R. Thus, our computational pipeline of type-II functional divergence between duplicate clusters can be used to reduce unexpected side effects and enhance the selectivity of therapeutic drugs.

Sequence conservation is a powerful indicator of functional importance [42]. Functionally important residues are correlated with structurally important residues, which play roles in ligand (small molecule, peptide, agonist, and antagonist) binding and protein–protein interactions [42]. Thus, ligand binding sites are strongly related to sequence conservation. When drugs that typically interact with conserved residues exhibit drug promiscuity, either type-II or type-I functional divergence can be analyzed between paralogs to achieve targetable differences. We also confirmed the role of residues related to type-I functional divergence in the binding of ligand and agonists to GLP-1R. We computed the coefficient of type-I functional divergence (denoted by θ_I_) between GCGR and GLP-1R. The coefficient was θ_I_ = 0.4902 ± 0.1072, which was significantly larger than 0 and indicated the occurrence of type-I functional divergence between two paralogs. We identified a type-I-related residue Glu294 in the binding sites of GLP-1R. Glu294 is a functionally important site for the signaling mechanism and receptor activation [36], and exhibits high conservation in GLP-1R but showed variation at paralogous sites of GCGR. A typical pattern of type-I functional divergence was observed, i.e., conserved amino acids in one cluster and diverse amino acids in the other. In this study, we distinguished two paralogs based on type-I functional divergence and achieved tighter specificity control of drugs.

As binding sites are typically structurally important, such as the large conserved N-terminal extracellular domain for ligands binding in secretin-type GPCRs [43], we predicted that these sites are conserved in sequence even in different paralogs. GCGR antagonist antibodies mAb1, mAb23, and mAb7 target the ligand-binding cleft in the extracellular domain. Our sequence conservation analysis of these antagonists illustrated that most binding-site residues showed good conservation between paralogous GCGR and GLP-1R (chi-square test showed statistically significant p-values of 0.0003, 0.02, and 0.002 in mAb1, mAb23, and mAb7, respectively). There are also some variable residues other than type-II-specific residues in the binding sites. There may be some underlying mechanisms involving variable residues in the discrimination of GCGR and GLP-1R. For example, mutations in these variable residues showed structural differences such as a shift or changes in orientation of some side chain residues; thus resulting in some reduction in receptor activation and prevention of ligand binding [44]. However, this was not discussed in our study. Because most binding sites exhibit sequence conservation, type-II functional divergence residues are critical determinants for the selective binding of drugs to targetable receptors, particularly when there are no variable residues in binding sites. These findings may have important implications in the design of drug binding sites and reduction of off-target effects. Our results provide a foundation for improving efficiency and reducing costs in the rational design of drugs.

## Acknowledgements

This work was supported by a grant from the National Science Foundation of China (31571355, 31301034). G.X. was supported by grants from Fudan University and Iowa State University.

